# Genome-scale sequence disruption following biolistic transformation in rice and maize

**DOI:** 10.1101/379644

**Authors:** Jianing Liu, Natalie J. Nannas, Fang-fang Fu, Jinghua Shi, Brooke Aspinwall, Wayne A. Parrott, R. Kelly Dawe

## Abstract

We biolistically transformed linear 48 kb phage lambda and two different circular plasmids into rice and maize and analyzed the results by whole genome sequencing and optical mapping. While some transgenic events showed simple insertions, others showed extreme genome damage in the form of chromosome truncations, large deletions, partial trisomy, and evidence of chromothripsis and breakage-fusion bridge cycling. Several transgenic events contained megabase-scale arrays of introduced DNA mixed with genomic fragments assembled by non-homologous or microhomology-mediated joining. Damaged regions of the genome, assayed by the presence of small fragments displaced elsewhere, were often repaired without a trace, presumably by homology-dependent repair (HDR). The results suggest a model whereby successful biolistic transformation relies on a combination of end joining to insert foreign DNA and HDR to repair collateral damage caused by the microprojectiles. The differing levels of genome damage observed among transgenic events may reflect the stage of the cell cycle and the availability of templates for HDR.

## INTRODUCTION

The creation of genetically modified crop lines through transformation is typically performed using *Agrobacterium-mediated* gene transfer ^1^ or particle bombardment ^2^. Both modes of transformation insert recombinant DNA in a random and uncontrolled manner. *Agrobacterium* is viewed as superior because it often delivers complete gene constructs bounded by known left and right borders ^1^. The integration of *Agrobacterium* transfer DNA (T-DNA) occurs at spurious double strand breaks through the activity of native polymerase theta and microhomology-mediated repair ^3^. Despite its relative precision, most T-DNA insertions are at least dimers ^3^ and many are composed of long arrayed multimers ^4–6^. In addition, *Agrobacterium* transformation may result in multiple T-DNA insertions across the genome, and integration can result in large deletions ^7,8^ as well as chromosomal inversions, translocations, and duplications ^4,8–12^.

Biolistic transformation offers the advantage that it can deliver any form of DNA, RNA or protein ^13–16^, a property that has been exploited to facilitate gene editing technologies ^17–20^. Biolistic transformation is also free of the constraints associated with *Agrobacterium-host* plant interactions. Unaltered BAC sequences larger than 100 kb ^21,22^ and an intact linear 53 kb molecule ^23^ have been integrated into plants by biolistic methods. Similarly, very long PCR products containing >100 kb of a simple repeating structure (ABS arrays) were co-bombarded with a selectable marker plasmid to create maize transgenics with inserts ranging from ~200 to 1000 kb in size ^24^ However, transgene copy number following biolistic transformation can be very high (depending on the amount of DNA delivered into cells ^13^) and very little is known about the process or mechanism of insertion following biolistic transformation. Prior literature based primarily on Southern blots indicates that sequence breakage and reassembly is common ^25–29^. The only detailed sequence-level analysis of transgenes following biolistic transformation revealed a few large fragments and many small shattered pieces, with 50 of 82 insertions being less than 200 bp in length ^27^. These limited sequence data suggest there may be unexpected and severe genomic consequences associated with biolistic transformation.

As a means to better understand the mechanistic underpinnings of biolistic transformation, we transformed linear and circular DNA molecules into rice (*Oryza sativa*) and maize (*Zea mays sp mays*) and subjected the lines to whole genome sequencing and analysis. The data revealed a wide spectrum of insertions and outcomes, from simple insertions to extraordinarily long shattered arrays. Multiple forms of genome damage were observed, including chromosome breakage and shattering and extreme copy number variation. We also found evidence of homology-directed repair (HDR) at sites that had been damaged during transformation. The data indicate that transformation involves both damage to the genome and fragmentation of the input DNA, creating tens to thousands of double stranded breaks that are repaired by end-joining and HDR in ways that can either create simple insertions or cause large structural changes in the genome.

## RESULTS

### General assessments of the genomes after co-bombardment with lambda and plasmid

We biolistically transformed 48 kb linear lambda phage DNA ^30^ and appropriate selectable marker plasmids into rice and maize using a 2-fold (rice) or 4-fold (maize) molar excess of lambda. All sequence analyses were carried out on genomic DNA from cultured callus tissue to obtain an unvarnished view of the transformation process; however three of the rice lines and all of the maize lines were also regenerated to plants (Supplementary Table 1). After screening the transformed callus by PCR to confirm the presence of lambda, we sequenced 14 rice lines and 10 maize lines at low coverage. The data revealed that over a third of the rice events contained less than one copy of lambda while the remaining two-thirds contained approximately 1 to 43 copies (Supplementary Table 1, where copy number is a sequence coverage value, and does not imply that any single lambda is intact). The maize transgenic events showed a similar wide range from approximately 1 to 51 copies (Supplementary Table 1). The selectable marker plasmids were observed at lower abundances reflecting their lower representations during transformation.

To interpret the distribution and structure of the insertions, eight rice lines and four maize lines were sequenced at 20X coverage by 75 bp paired-end Illumina sequencing (Table 1). The data were then interpreted using SVDetect, which employs discordant read pairs to predict breakpoint signatures through clustering ^31^, and Lumpy, which uses discordant read pairs and split reads to determine SV types by integrating the probabilities of breakpoint positions ^32^. The paired end Illumina reads were aligned to the rice or maize reference genomes with the complete lambda and plasmid sequences concatenated as separate chromosomes. Insertions were detected as inter-chromosomal translocations between lambda, plasmid and genome, whereas rearrangements were identified as intra-chromosomal translocations. Based on simulations using in silico modified forms of rice chromosome 1 with randomly inserted lambda/chromosomal fragments, we estimate that our approach identifies about 84% of the breakpoints involving lambda and about 66.5% of the junctions involving two chromosomes but no lambda (Supplementary Table 2).

**Table 1.** Copy number of introduced molecules (lambda and co-bombarded plasmid) and number of breakpoints in rice/maize transgenic genome

| Transgenic events† | Genome coverage | Copy Number |  | Number of breakpoints |  |  |  |  |  |
| --- | --- | --- | --- | --- | --- | --- | --- | --- | --- |
|  |  | Lambda | Plasmid | Lambda-Lambda | Lambda-plasmid | Lambda-genome | Plasmid-genome | Intra-genome | Inter-genome |
| Os λ-1 | 22.45 | 0.12 | 3.63 | 1 | 11 | 0 | 2 | 2 | 1 |
| Os λ-2 | 20.95 | 0.95 | 0.80 | 19 | 1 | 3 | 0 | 2 | 0 |
| Os λ-3 | 20.84 | 0.04 | 1.37 | 0 | 3 | 1 | 2 | 1 | 1 |
| Os λ-4 | 19.63 | 32.48 | 3.87 | 517 | 21 | 14 | 0 | 1 | 0 |
| Os λ-5 | 21.58 | 37.06 | 2.22 | 420 | 18 | 18 | 0 | 1 | 0 |
| Os λ-6 | 21.78 | 17.35 | 1.18 | 257 | 6 | 123 | 0 | 1 | 13 |
| Os λ-7 | 19.64 | 1.72 | 1.07 | 51 | 4 | 63 | 3 | 12 | 14 |
| Os λ-8 | 21.84 | 7.83 | 0.98 | 152 | 3 | 99 | 1 | 67 | 40 |
| Zm λ-1 | 14.07 | 1.73 | 1.88 | 22 | 22 | 17 | 6 | 1 | 0 |
| Zm λ-2 | 18.05 | 19.11 | 13.97 | 31 | 30 | 15 | 5 | 14 | 19 |
| Zm λ-3 | 15.55 | 10.33 | 2.00 | 12 | 8 | 8 | 1 | 1 | 8 |
| Zm λ-4 | 17.93 | 40.48 | 20.50 | 241 | 143 | 73 | 4 | 10 | 18 |

† Transgenic events of rice (*Oryza sativa*) and Maize (*Zea mays*) are labeled as “Os” and “Zm” respectively.

The sequence data also allowed us to identify deletions and duplications of genomic DNA by changes in read depth as assayed by CNVnator ^33^. Unique breakpoints and regions showing copy number variation were plotted using the Circos chromosome visualization software (Fig. 1, 3, 4 and Supplementary Fig. 1 and 3). We found a wide range of sequence complexity, ranging from simple insertions to long complex arrays and massive genome-scale disruptions.

**Fig. 1.**
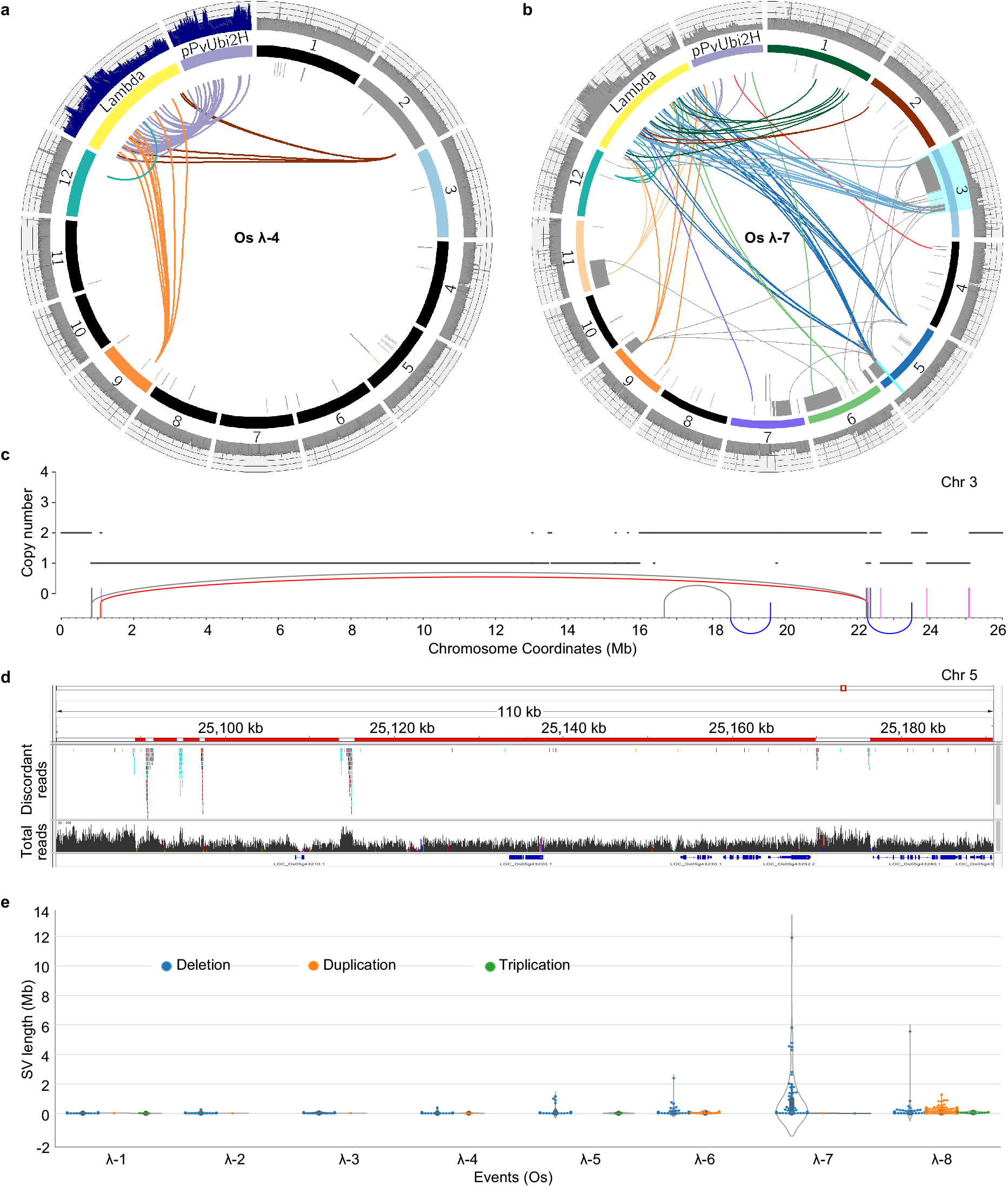
Spectrum of genomic outcomes following transformation with lambda and plasmid in rice. All Circos plots are annotated as follows. The twelve rice chromosomes are shown along with λ and plasmid pPvUbi2H magnified at 1,000X and 5,000X. The outer track shows sequence coverage over each molecule or chromosome as histograms. The inner ring demonstrates DNA copy number profiles derived from read depth, with grey shown as 1 copy, orange as 3 copies, dark red as 4 copies and black as more than 4 copies. The inner arcs designate inter- and intra-chromosomal rearrangements. Breakpoints within the genome are colored grey while the breakpoints between λ or plasmid and the genome are colored to match the respective chromosomes. **a**, Rice event λ-4, which contains a long transgene array in chromosome 2. The coverage in histogram tracks of λ and plasmid are divided by 15 and 1.5 respectively. **b**, Rice event λ-7, illustrating a complex event with severe genome damage. **c**, Two regions of rice λ-7 at high resolution. The horizontal lines show copy number states over the 26 Mb region on chromosome 3 highlighted in cyan. Vertical bars represent inter-chromosomal breakpoints (grey) and breakpoints involving λ (plum). The arcing links show local rearrangements of the deletion-type (grey), duplication-type (red), and intra-chromosomal translocation-type (blue). For a visual depiction of how local rearrangements are defined using paired end reads, see Supplementary Fig. 2. **d**, A region from 25.1 Mb to 25.2 Mb on chromosome 5 as visualized with IGV. Deleted regions are shown in red and retained regions in white (upper), as indicated by the alignment of discordant reads (middle) and read depth (lower). **e**, Swarm and violin-plots showing the distribution of the size and number of deletions, duplications and triplications in all rice events transformed with λ. Each dot in the swarm plots represents a different SV. Violin plots represent the statistical distribution, where the width shows the probability of given SV lengths.

### Simple low-copy insertions

Four rice lines and two maize lines had one or few insertions and otherwise did not show evidence of genome damage (Fig. 1, Supplementary Fig. 1 and 3). In these events, there were fewer than 40 detected breakpoints between lambda, plasmid and chromosomes (Table 1), and there were small chromosomal deletions of less than 20 kb around insertion sites (Fig. 1, Supplementary Fig. 1 and 3). For example, in rice λ-1 there is a 27 kb insertion composed of rearranged lambda (5.8 kb) and plasmid (21.2 kb) fragments in a region of chromosome 8 that has sustained an 18 kb deletion. Similarly, maize λ-1 contains 86.3 kb of combined lambda and plasmid DNA in chromosome 9 with no deletion at the point of insertion, and rice λ-4 (discussed in detail below) contains a long array of lambda and plasmid fragments in chromosome 2 and a small 9 bp deletion at the site of insertion. In these and other cases of simple insertions, there was no other evidence of chromosome truncation or duplication as judged by read depth.

### Creation of long arrays

Several transformants had large amounts of lambda DNA. Rice λ-4 is the simplest of these, with lambda junctions involving three genomic locations (chromosome 2, 9 and 12), and no other evidence of genome damage (Fig. 1a, 1e). As assayed by sequence coverage and SV estimates, this event contains the equivalent of 37 copies of lambda broken into a minimum of 552 pieces. Local sequence assembly indicated that the apparent insertions in chromosome 9 and 12 are small sections of chromosomal DNA flanked on both sides by lambda fragments. Two fragments of chromosome 9 are 102 and 464 bp in length, and one fragment of chromosome 12 is 108 bp in length (see example in Fig. 2b, Supplementary Fig. 4a). In contrast, on chromosome 2, the assemblies revealed two simple lambda-genome junctions. These data suggest that rice λ-4 has a large insertion on chromosome 2 and that small sections of chromosome 9 and 12 are intermingled within it. Analysis of 23 self-cross progeny from rice λ-4 supported this view showing that the fragments of chromosome 9 and 12 and the junctions on chromosome 2 are genetically linked (Supplementary Fig. 4b).

**Fig. 2.**
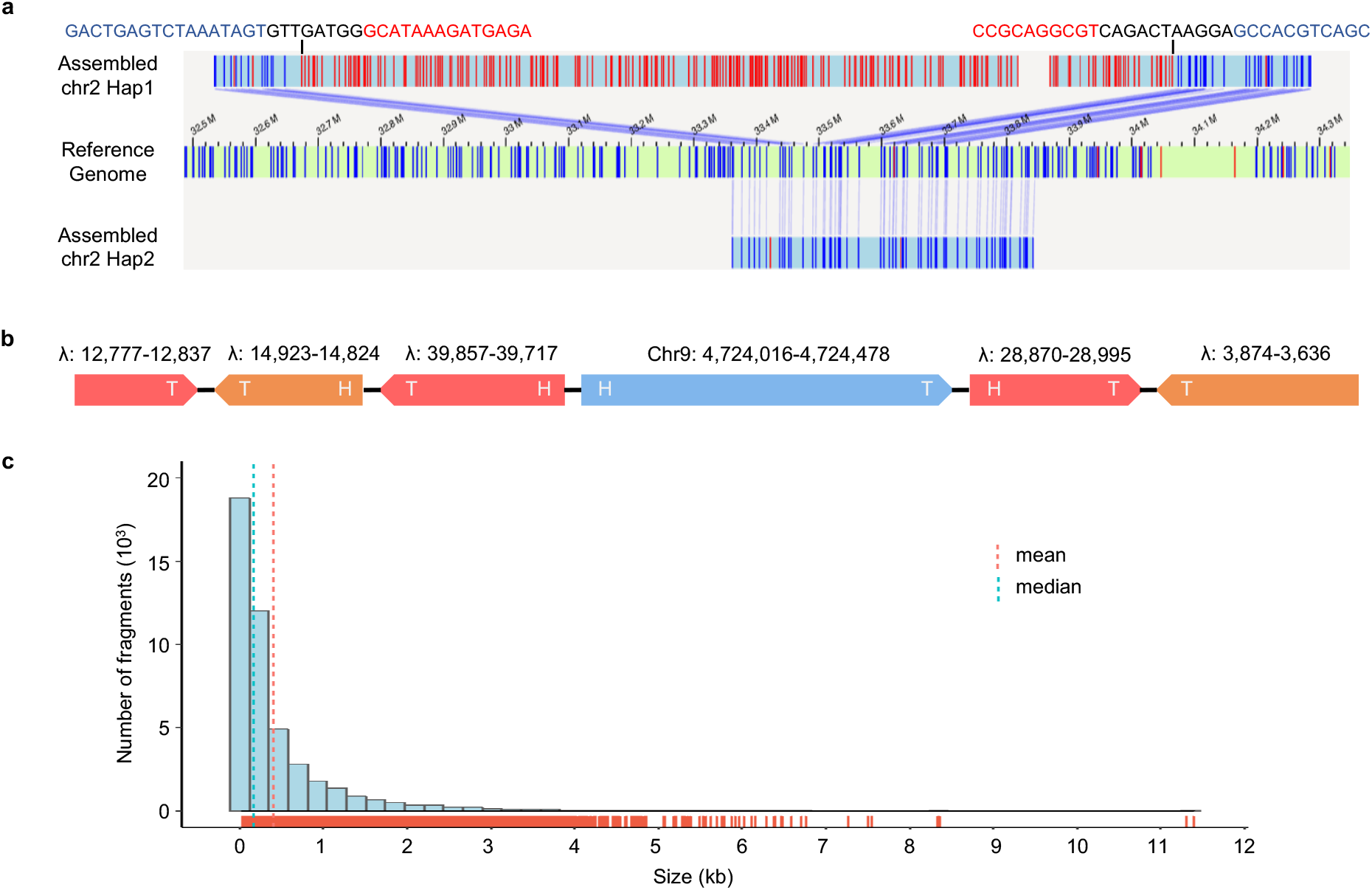
Characteristics of the long transgene array in rice event λ-4. **a**, Bionano assembly depicting the 1.6 Mb insertion in chromosome 2. The middle panel represents the reference genome, and the upper and lower panels depict the assembled transgenic and wild type chromosomes in this heterozygous line. The blue bars indicate matching restrictions sites between the reference and assembled contigs, and red bars denote restriction sites within the insertion. The nucleotide sequences above the upper panel shows the breakpoint sequences, with chromosome sequence highlighted in blue, λ sequence highlighted in red, and novel sequences in black. **b**, A 1.1 kb region assembled from Illumina data showing five λ pieces and a single fragment of chromosome 9 in rice event λ-4. The direction of the arrows indicate the 3’ ends (Tails) of λ and chromosomal genomic fragments. Four different orientations between intra- and inter-chromosomal pieces can be found in this sequence: Tail (3’)-Head (5’), Tail-Tail, Head-Tail, Head-Head. **c**, Size distribution of λ fragments in the array as determined by PacBio sequencing.

To confirm our interpretation of the rice λ-4 event, we analyzed the original T0 plant by Bionano optical mapping, where long DNA molecules were fluorescently labeled at the restriction sites BspQI, imaged, and assembled into megabase-scale restriction map contigs ^34^. The data revealed no insertions on chromosome 9 or 12, but an unequivocal large insertion on chromosome 2 at the location predicted. There are two assemblies over this region, one for the wild type chromosome 2 and one showing a large insertion of at least 1.6 Mb containing novel sequence. The 48-kb lambda molecule contains six BspQI sites in a distinctive pattern. However, Bionano alignment software failed to detect any similarity between lambda and the BspQI recognition pattern within the array on chromosome 2, as expected if lambda molecules were broken and rearranged. To more accurately assess the internal structure of the array, we sequenced a T1 plant that was homozygous for the insertion on chromosome 2 at 25X coverage using PacBio technology. A total of 1810 (45280/25) lambda fragments ranging in size from 31 to 11387 bp were identified. Over 96% of the lambda fragments were less than 2kb with a mean fragment size of 410 bp (Fig. 2c).

Evaluation of breakpoint junctions provided information on the mechanisms of repair that operate to create long arrays (Supplementary Fig. 5a). The two major forms of non-homologous repair are nonhomologous end joining (NHEJ), which is typified by blunt end junctions and short insertions ^35^, and microhomology-mediated end joining (MMEJ), which is characterized by junctional microhomology of at least 5 bp ^36^. Computational analyses of the junctions in rice and maize transgenic events revealed blunt-end connections (25%), short insertions varying in size from 1 to 80 bp (21%) and junctions displaying microhomology in the range of 1-4 nucleotides (50%) and 5-25 nucleotides (4%), suggestive of both NHEJ and MMEJ (Supplementary Fig. 5b). Further, the four relative orientations of lambda fragments (tail-head, tail-tail, head-tail, and head-head), were nearly uniformly distributed (Supplementary Fig. 5c) as expected for a random rejoining process. We also investigated whether the natural overlapping single stranded ends of lambda (the 12-bp cos sites ^30^) may have played a role in multimerization. The data showed that five rice lines and three maize lines contained a single annealed cos site; a low frequency that supports the view that homology-based annealing and ligation have minor roles in the assembly of broken fragments.

### Evidence of HDR

A second major form of repair is homology-directed repair (HDR) where double stranded breaks are seamlessly corrected using undamaged homologous molecules such as sister chromatids as templates. If a segment of the genome is broken away and not repaired, we expect to find a deletion at the original coordinates, whereas if the damaged region is repaired by HDR, we expect to find no evidence of damage at the original coordinates. Incorporation of a displaced fragment at a new location followed by repair of the original site will result in a total of three copies of the region affected.

The analysis of rice λ-4 revealed that small sections of chromosome 9 and 12 were included in a long array of lambda fragments but that there were no changes from wild type where the original damage occurred (as assayed by optical mapping, Fig. 2a). We also analyzed the coordinates surrounding the affected sites on chromosomes 9 and 12 (plus or minus 1 Mb) for a clustering of discordant reads or significant changes in read depth and found no evidence of sequence disruption. Further, PCR analysis of the T0 line revealed no evidence of small deletions at these coordinates. These data are consistent with a model where chromosome 2 and 9 were damaged, broken fragments were included in the assembly of the long chromosome 2 array, and the damaged chromatids were repaired by HDR.

To determine if HDR had occurred in any of the other lines assayed, we identified 78 additional displaced genomic fragments in 4 rice events and 4 maize events. We then systematically checked for increases in read depth and clusters of discordant reads that map to the native locations of these displaced fragments. The data provide evidence of HDR in 3 rice events and 3 maize events (Supplementary Table 3, 4). For example, a 110 bp displaced fragment from chromosome 1 and a 69-bp fragment from chromosome 9 in rice λ-5, both flanked by lambda pieces, exhibited increased sequence coverage by 50% and no apparent deletions at the original coordinates (Fig. 3a). While most of the displaced genomic fragments in lambda arrays were on the order of a few hundred bases, we also found evidence of breakage and repair among the chromosomes on a larger scale (Supplementary Table 3, 4). For example in rice λ-8, a 21 kb and a 34 kb region from chromosome 2 were broken away, connected by a small fragment from lambda and reinserted in the genome, followed by repair at the original locations. This resulted in duplication regions clearly visible by read depth (Fig. 3b). The limits on the size of a deletion that can be repaired by HDR are not known, but in animals HDR can be used to incorporate new (knock-in) constructs as large as 34 kb ^37^.

**Fig. 3.**
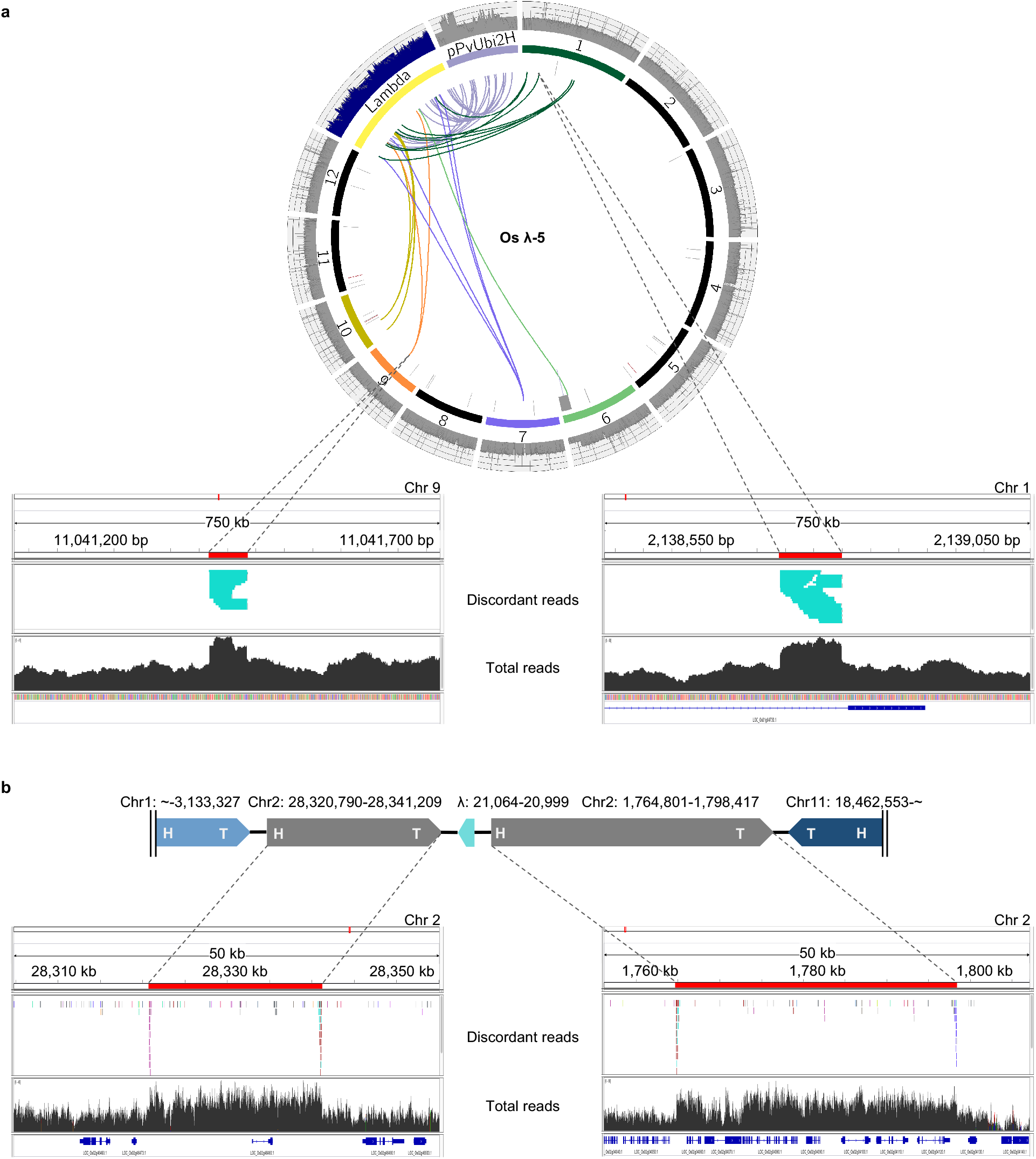
Evidence of HDR in rice transgenic events. **a**, Circos plots of rice transgenic event λ-5 annotated as in Figure 1. The λ coverage is divided by 15. Region 2,138,442 - 2,139,257 on chromosome 1 and region 11,041,419 - 11,041,484 on chromosome 9 are displayed in IGV windows, where displaced fragments (110 bp and 66 bp) are highlighted in red. The upper panels shows only discordant reads (where one end maps to the fragment and the other maps to another chromosome). The lower panels show all reads, illustrating the ~50% increase in read depth indicative of an HDR event. **b**, Complex rearrangements observed in rice event λ-8. Regions from chromosome 2 were assembled into an array with other broken fragments at an unknown location in the genome. The damaged regions of chromosome 2 were subsequently repaired as demonstrated by the ~50% increase in read depth.

### Deletions and evidence of breakage-fusion-bridge cycling

Copy number profiling provided evidence for many deletions ranging in size from 3.5 kb to 11.9 Mb in rice and 115 kb to 62 Mb in maize (Supplementary Table 6). Deletions and duplications/triplications greater than 1 Mb were found in four rice events and three maize events (Fig. 1e, Supplementary Fig. 3b, Supplementary Table 6). Deletions were particularly common around transgene insertions and at the ends of chromosomes, and the majority were associated with the presence of lambda or plasmid DNA, indicating that the breaks occurred as a consequence of the transformation process. Deletions that appeared to have no connection with lambda or plasmid may either reflect our imperfect (84%) ability to detect such junctions, or identify regions that were damaged and repaired without the involvement of introduced DNA. No deletions were observed in the single non-transformed rice callus line used as a control.

Chromosome breakage is expected to yield a double stranded break that is repaired by ligation to an introduced DNA molecule or to another broken chromosome. The fusion of two different chromosomes can cause the formation of a dicentric chromosome that is unstable during mitosis. When the centromeres on a dicentric chromosome move in opposite directions during anaphase, the pulling forces cause a re-breaking of the chromosome that initiates a breakage-fusion-bridge (BFB) cycle that may repeat for many cell divisions ^38–40^. The BFB cycle can lead to local duplications and higher order expansions ^41,42^. Chromosomes 4 and 7 in rice λ-8 showed evidence of BFB, with over half of a chromosome amplified in a region displaying inversions (Fig. 4a). These chromosomes appear to be partially trisomic, and may be the result of inactivation of one of the two centromeres. Copy number gains were also observed on chromosome 6 in maize λ-4, where the amplified regions are adjacent to a terminal deletion (Fig. 4d). At least two inversions of 3.9 Mb and 2.8 Mb were found in the amplified area. Read depth increases adjacent to a terminal deletion were also found on chromosome 9 in maize λ-3, where the amplified region displayed switches from 2 to 6 copies (Fig. 4c).

**Fig. 4.**
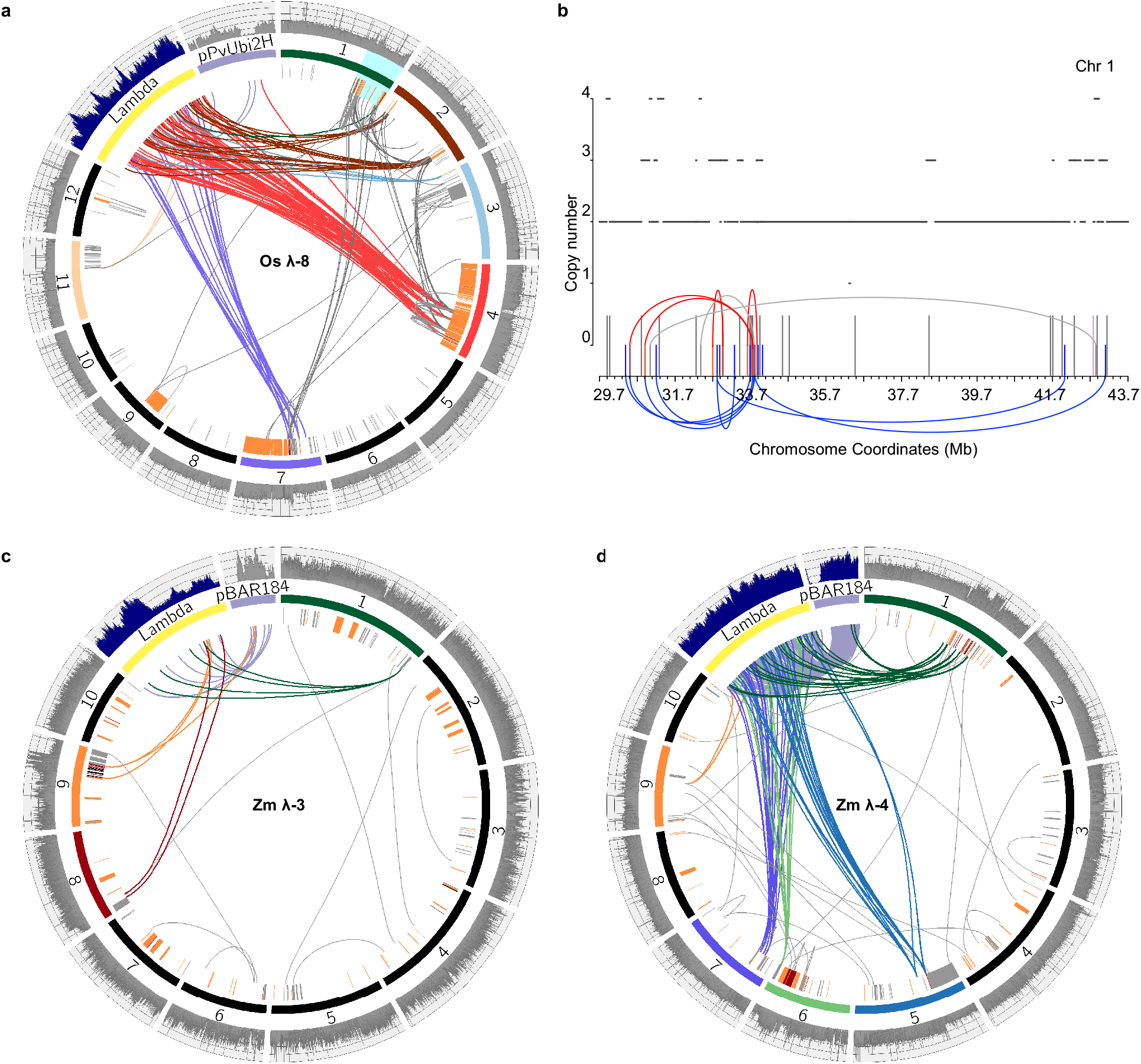
Chromothripsis-like outcomes and BFB (breakage-fusion-bridge)-like genomic rearrangements in rice and maize transgenic events. **a**, Circos plot of rice transgenic event λ-8 annotated as in Figure 1. The coverage of λ in the histogram track is adjusted to 4X. **b**, Copy number states of region 29.7 - 43.7 Mb on chromosome 1 (highlighted in cyan) are shown on the right and annotated as in Figure 1c. **c**, Circos plot of maize transgenic event λ-3, with coverage of λ in the histogram track divided by 5. Note the region of increased copy number states on chromosome 9 indicative of BFB. **d**, Circos plot of maize transgenic event λ-4, with the coverage of λ and plasmid in the histogram track divided by 15 and 10 respectively. Note the regions of increased copy number states on chromosome 1 and 6 indicative of BFB.

### Shattering and chromothripsis-like outcomes

Animal cells sustaining chromosome loss or breakage undergo a process known as chromothripsis that results in complex genomic rearrangements in localized areas, generally consisting of tens to hundreds of small pieces ^43,44^. The reassembly process involves a reshuffling and loss of sections of the genome. Instead of uniform coverage, a region that has undergone chromothripsis shows oscillations from the normal copy number state of two to a copy number state of one (haploid) and occasionally three (triploid) in the context of numerous rearrangements. Analysis of the rice and maize transgenic events revealed similar oscillating copy number states in regions surrounding what appear as “impact sites” on Circos displays: large areas of genome damage with multiple lambda and plasmid fragments.

We found particularly complex rearrangements with copy number oscillations and interspersed lambda and plasmid fragments in three rice events. In rice λ-7, broken fragments (44bp to 7858bp) from localized regions of chromosome 3, 5, 6, 7, 9 and 11 were interlinked along with lambda and plasmid fragments in inverted and non-inverted orientations (Fig. 1b, 1c, 1d). These patchwork assemblages are presumably integrated into one or a few arrays. The damage imparted during transformation caused large swathes of the same regions on chromosomes 3, 5, 6, 7, 9 and 11 to be deleted. The combination of retained displaced fragments and deletions results in oscillating patterns between 1 and 2 copy number states (Fig. 1c, 1d). Higher order oscillation patterns were identified in rice λ-8, where numerous fragments from chromosome 1 were linked with segments of chromosome 2, 4, 7, 9 and 11 in what is likely another complex array (Fig. 4a). However in this case the read depth data indicate that the damaged regions of chromosome 1 were repaired by HDR. The combination of retained displaced fragments and repaired regions result in oscillating patterns between 2 and 3 copy number states (Fig. 4b).

Similar results were found in three maize events where large deletions and duplications occurred. The sensitivity of our assay is significantly lower in maize because of the high repeat content and necessity of using only perfectly mapped reads. Although we can only detect a fraction of the rearrangements present, the linking patterns between displaced genomic segments and lambda and plasmid is obvious (Fig. 4c, 4d and Supplementary Fig. 3a). For example, maize λ-4 shows lambda and plasmid within an inter-chromosomal network including sections of chromosomes 1, 5, 6, 7, and 9, as well as evidence of copy number switching (Fig. 4d).

### Similar genome scale disturbances in single plasmid transformations

We were concerned that the linearity of lambda or the high concentration of DNA used when transforming lambda may have led to new or extreme forms of genome damage. To test whether this was the case, we transformed rice with circular plasmids designed to knockdown (pANIC10A-0sFPGS1) or overexpress (pANIC12A-OsFPGS1) folylpolyglutamate synthetase 1 (chosen for its presumptive role in regulating lignin content). The first plasmid (“10A”) is 17603bp and the second (“12A”) is 17501bp. Approximately 125 ng of DNA was delivered to 100 mg of callus tissue per shot, which is considerably lower than the 585 ng of DNA delivered for lambda. In addition we only sequenced the genomes of fully regenerated plants.

The rice lines transformed with single plasmids showed a narrower span of transgene copy numbers (ranging from 0.5 to 12.3X, Supplementary Table 5) and the number of breakpoints between plasmid and chromosomes was considerably lower, consistent with the smaller size of the input molecules and the lower amount of DNA used in transformation ^45^. The genome-level damage was not as extensive as what we observed for the lambda transformation experiments. However, as in the lambda experiments, there were large-scale deletions, inversions, duplications consistent with BFB, and rearrangement patterns indicative of chromothripsis-like processes (Fig. 5, Supplementary Fig. 6 and 7). For example, in event 12A-6, chromosome 4 sustained a large deletion and the remainder of the chromosome was duplicated to create a region of partial trisomy (Fig. 5d). Evidence of alternating copy number states was found on chromosome 1 in event 10A-6 (Fig. 5e), and chromosome 8 in event 12A-3 (Supplementary Fig. 7b).

**Fig. 5.**
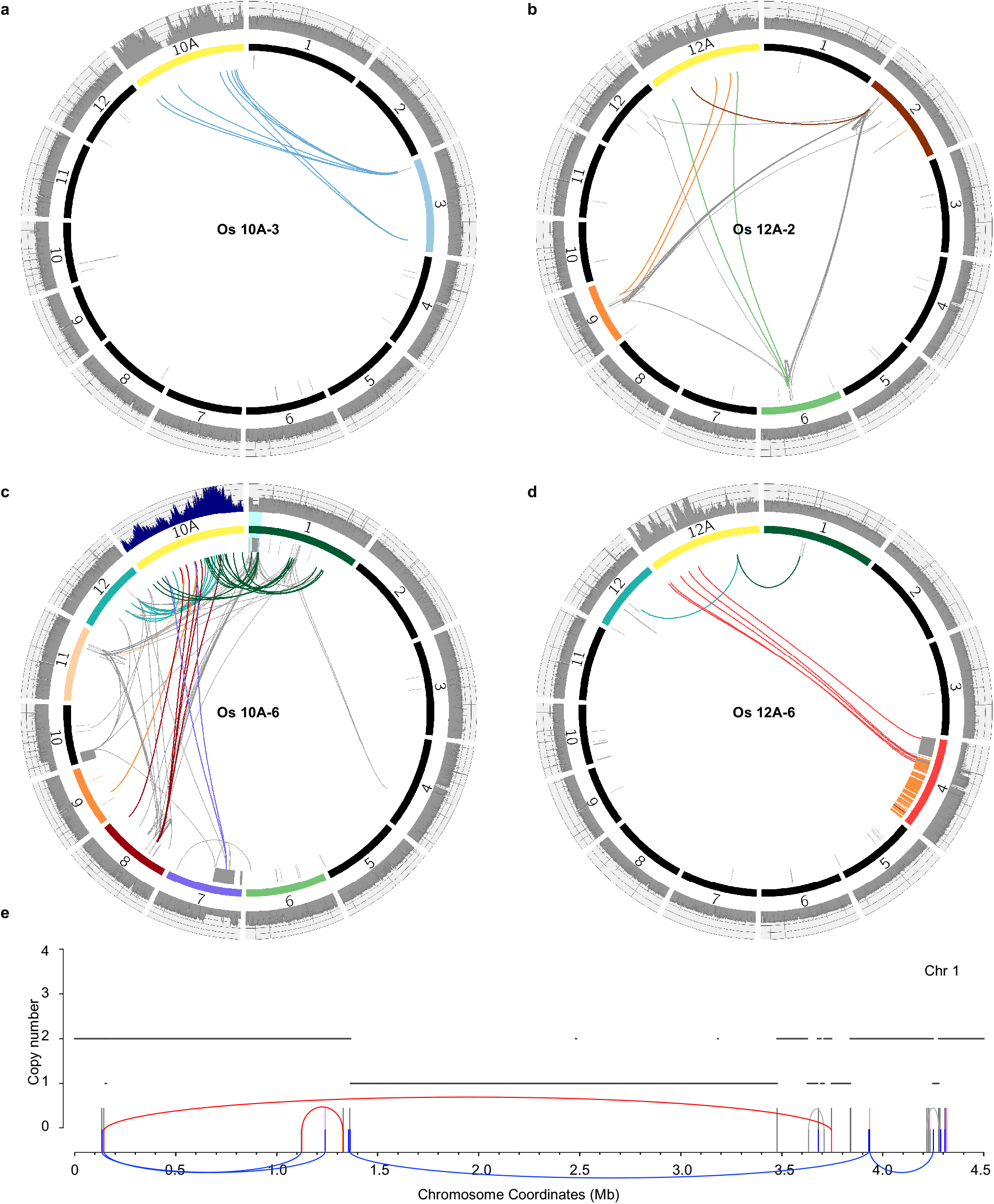
Similar genomic disturbances following single plasmid transformations. Circos plots of rice lines transformed with plasmid pANIC10A-OsFPGS1 (**a,c**) and pANIC12A-OsFPGS1 (**b,d**). **a**, Simple insertion. **b**, Complex insertion showing a network of interlinked genomic regions. **c**, Extensive damage with a deletion on chromosome 7 and apparent chromothripsis on chromosome 4 (coverage of the 10A plasmid is divided by 2). **d**, Chromosome-scale disruption with a partially trisomic chromosome 4. **e**, Copy number states of the chromothripsis region of chromosome 1 of “c” (region 0 - 4.5 Mb, highlighted in cyan) are shown on the lower panel and annotated as shown in Figure 1c.

## DISCUSSION

Here we provide new data showing that biolistically transformed rice and maize plants contain a wide diversity of transgene copy numbers ranging from a fraction of a single copy to as many as 51. The results are consistent with prior data from rice, showing that biolistic methods tend to deliver more transgene copies than Agrobacterium ^46^, as well as maize, where one study showed that the majority of biolistically transformed lines contained more than 10 transgene copies and several events contained more than 100 copies ^25^. We also join multiple prior authors in providing evidence for long arrays of broken and rearranged plasmids ^26,47–50^.

Key among the early studies was work from the Somers lab ^26,27^ showing that plasmids transformed biolistically are frequently broken into small (<100 bp) pieces and scrambled with genomic segments to produce very complex insertions. Based on the analysis of 155 breakpoints ^27^ the authors speculated that DNA was broken randomly and rejoined at blunt ends often containing microhomology ^27^. Our more extensive analysis implicates non-homologous end joining (NHEJ) as the primary mechanism for rejoining broken fragments and that microhomology-mediated end joining (MMEJ) is involved as a secondary pathway.

Traditional transformation involves transforming explanted plant tissues, or dedifferentiated callus tissue obtained from them, and then treating the tissue with hormones that promote plant regeneration. Callus is known to tolerate chromosome instability ^51^ and we might expect to see more severe genome damage at this stage. In contrast, organogenesis and embryogenesis require complex developmental programs that may select against severe genome imbalance. By sequencing the lambda lines at the callus stage, we may have identified forms of genome damage than would not have been present had we sequenced regenerated plants. Likewise, the fact that we observed less genome damage using single plasmids may be partially attributable to the fact that the plants had been regenerated. Nevertheless our data from two species using different DNA inputs, and assayed pre- or post-regeneration, indicate that biolistic transformation can cause significant genome-scale disruption. We find that large deletions, inversions, duplications, genome shattering, and sites of homology-directed repair are frequently associated with fragments of introduced DNA. The intermingling of sequence suggests that the mutations were a result of transformation and presumably a direct outcome of the impact of microprojectiles. Such damage is to be expected, as the 0.6 μm gold beads used for biolistic transformation are about a third the diameter of a rice nucleus (about 2 μm ^52^) and 300 times larger than the diameter of DNA. When the genome is damaged in this manner, it can be repaired in one of three ways (Fig. 6):

1. Repair can occur by homology-directed repair such that the damaged region is completely restored to its original state (Fig. 6d).
2. Repair can occur by NHEJ or MMEJ, where the end of any other broken DNA molecule is used as a substrate. Broken fragments of introduced DNA are a likely substrate particularly when they contain markers that are under selection. The other end of the newly joined fragment may then be ligated to a second fragment of introduced DNA or to another segment of the genome. If this process culminates by reconnecting the two pieces of the original chromosome, the result will be a “simple insertion” containing a variable number of conjoined foreign DNA fragments (Fig. 6a).
3. Repair can be initiated by the process above but not culminate in the reconstitution of the original chromosome. The break may not be repaired at all or it may culminate in connecting of two different chromosomes. In this case there can be severe genomic consequences including large terminal deficiencies, chromosome fusion and BFB cycling, and more complex events resembling chromothripsis (Fig. 6b, 6c). These dramatic chromosomal rearrangements are a natural outcome of the same processes that are used to create a simple insertion.

**Fig. 6.**
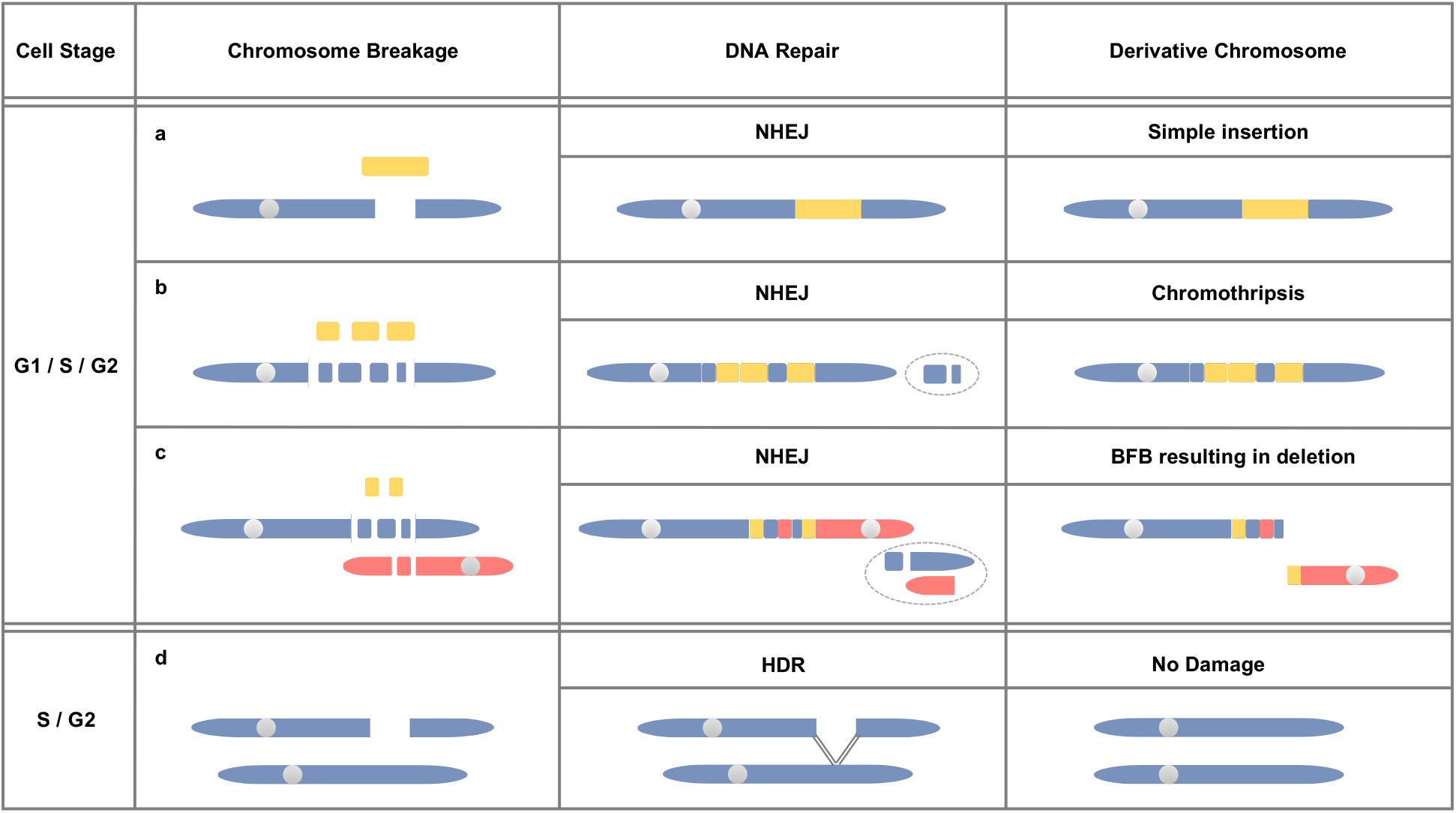
Models for genomic outcomes after biolistic transformation. The stage of cell cycle may influence the outcome of biolistic transformation. The models are based on the fact that in animals and presumably plants, non-homologous end joining (NHEJ) is the most likely repair pathway in G1 and homology directed repair (HDR) is more likely in S and G2. **a**, Simple insertion. Fragments of introduced molecules (yellow) are ligated with broken ends of native chromosomes by NHEJ (non-homologous end joining). **b**, Chromothripsis-like genome rearrangements. Localized regions from native genome are shattered resulting in many double stranded breaks. Fragments of chromosomes and introduced molecules are stitched together through NHEJ, creating complex patterns that involve the loss of genomic DNA and changes in copy number state (lost regions are circled). **c**, Breakage and joining of two different chromosomes and breakage-fusion-bridge (BFB)-like genome rearrangements. When two chromosomes are broken they can be ligated together through NHEJ. The resulting dicentric chromosome is expected to undergo BFB, which can result in stable terminal deletions. **d**, DNA damage repaired by HDR. Double stranded breaks in S or G2 phase may be repaired by HDR through recombination with an intact sister chromatid.

The stage of the cell cycle may have a significant impact on the outcome of biolistic transformation. Data from non-plant systems indicate that while NHEJ is active throughout interphase, it is particularly important in G1. In contrast, HDR is more likely in S and G2 phases after DNA replication has provided additional templates for repair ^53–55^. Simple insertions may be more probable when the cell is transformed in S or G2 so that NHEJ can insert the foreign DNA while HDR serves to repair extraneous damage. Simple insertions may also be an outcome of transformation during mitosis when chromosomes are distributed in the cytoplasm. DNA introduced during metaphase or anaphase might find its way into newly-forming telophase nuclei, and subsequently be inserted into the genome as consequence of routine DNA repair (similar to T-DNA ^3^).

When chromosomes are broken in G1, deletions and translocations are to be expected. We observed many examples of chromosomes that were missing large terminal segments of chromosome arms (Fig. 1b, 3a, 4c, 4d, and 5c, 5d). The formation of a stable truncated chromosome requires that the end be healed by formation of a telomere which is a process that occurs over a period of cell divisions ^56,57^. In the period when there is an unattended double strand break without a stable telomere, the break is likely be repaired by NHEJ using any other broken chromosome. As famously described by Barbara McClintock ^56^, the fusion of broken chromosomes can initiate a breakage-fusion-breakage (BFB) cycle and amplification of genome segments on the affected chromosomes. In several cases we observed copy number states of 4, 5 and 6 that are difficult to explain by any other mechanism. We also observed partial and fully trisomic chromosomes (Fig. 4a, 5d). Such large-scale chromosome abnormalities may also be the result of the tissue culture process itself ^51^, and we cannot rule out the possibility that some of the chromosomal changes were either present before transformation or occurred after transformation. However, for most of the large duplications and deletions we observed, there was either evidence of inserted foreign DNA or evidence that the lost DNA had been fragmentation and rejoined with foreign DNA.

In addition, our analyses revealed extreme shattering and chromothripsis-like outcomes in several lines. Chromothripsis was originally described as a process whereby “tens to hundreds of genomic rearrangements occur in a one-off cellular crisis ^44^.” Our data meet this definition in a descriptive sense but the biological underpinnings are presumably different. For cancer lines, the simplest model (as it relates to our study, for other models, see ^58^) requires that a chromosome be partitioned from the primary nucleus, generally as a result of an error in chromosome segregation that leaves it stranded in the cytoplasm ^59^. The resulting micronuclei show aberrant DNA replication ^59–61^ and appear to have fragmented chromatin ^61^. The partially degraded chromatin can then be reincorporated into the primary nucleus where it is evident as broken and reassociated fragments ^58^. Recent data indicate that when plants sustain errors in chromosome segregation, they too show evidence of chromosome fragmentation with oscillating copy number states confined to single chromosomes ^62^. In contrast, our biolistically transformed lines are not expected to undergo regular loss of chromosomes during cell division. It is possible that microprojectiles severely damage nuclei such that portions of the genome are released into the cytoplasm. Another plausible explanation is that acentric fragments formed during the repair process (Fig. 6b, 6c) are lost during anaphase, become partially degraded, and are reincorporated into a nucleus during a subsequent cell division.

Taken together our data help to explain the long nearly continuous arrays of 156 bp repeats we observed following biolistic transformation of PCR products in maize ^24^ At the time we were unable to determine whether the long PCR products had been transferred intact or were broken and reassembled in planta. Based on the data here is seems more likely that the PCR products were fragmented and reassembled by NHEJ to create the observed long arrays. The longest intact fragment of lambda we observed was 11.3 kb (Fig. 2c). Although constructs as long as 53 kb have been recovered with careful selection for low copy inserts ^23^ and DNA is sometimes inserted cleanly without rearrangements at the point of insertion (e.g. Os λ-3), such events are noteworthy for their rarity. The normal outcome is shattered input molecules and significant disruptions of the genome.

From a product development perspective, genomic rearrangements were initially considered to be a food/feed safety hazard ^63^. To put this hazard in perspective, Anderson et al ^12^ noted that the genomic rearrangements from *Agrobacterium-mediated* transformation were an order of magnitude lower than those created by fast-neutron mutagenesis. In turn, rearrangements from fast-neutron mutagenesis were an order of magnitude lower than the standing genomic structural variations in the cultivated soybean germplasm pool, all of which has a history of safe use. The frequency of rearrangements from biolistic transformation may be more comparable to that induced by fast neutrons. Regardless, as of yet, there is no evidence that a genomic rearrangement has compromised the safety of a plant used as food ^64^, though its agronomic performance can be compromised. Since poor agronomic performance is not tolerated in modern cultivars and hybrids, there is a rigorous selection process that eliminates deleterious mutations during the breeding process ^65^.

From a research perspective, such rearrangements may be acceptable in some cases, while in others it may be necessary to consider that undetected rearrangements could be influencing the phenotype. Gene editing applications are a special case where the intent is usually to make a single precise change. Although there is great appeal in directly introducing Cas9 ribonucleoproteins ^18^ and repair templates ^20^ for this purpose, our data suggest that there is strong likelihood that the delivery method itself will cause unintended genome damage. Until new transformation methods become available, the *Agrobacterium-based* methods that have been in regular use for decades ^1^ remain the superior alternative in terms of minimizing genome rearrangements.

## METHODS

### Rice transformation

Rice variety Taipei 309 was transformed as described previously ^22^. For the lambda experiments, we mixed 33 ng of pPvUbi2H ^66^ which confers hygromycin resistance and a two fold molar excess (552 ng) of purified lambda DNA cI857 (New England Biolabs #N3011S). After screening for lambda by PCR (forward primer 5’-GACTCTGCCGCCGTCATAAAATGG and reverse primer 5’-TCGGGAGATAGTAATTAGCATCCGCC) 14 callus lines were chosen for sequence analysis. Three of these callus lines were regenerated to mature rice plants (Supplementary Table 1).

The plasmids pANIC10A-0sFPGS1 and pANIC12A-OsFPGS1 are based on the pANIC backbone ^66^ with inserts designed to silence or overexpress folylpolyglutamate synthetase. In these experiments only plasmid DNA was used, delivering ~125 ng per shot. All 12 lines were regenerated to plants.

### Maize transformation

Biolistic transformation of the maize inbred Hi-II was performed by the Iowa State University Transformation Facility (Ames, IA) as previously described ^67^. To achieve a four molar excess of lambda DNA, we mixed 20 ng of plasmid pBAR184 which confers resistance to glyphosate ^67^ and 528 ng of purified lambda DNA cI857 (New England Biolabs #N3011S). Ten callus lines were screened for lambda by PCR (forward primer 5’-GACTCTGCCGCCGTCATAAAATGG and reverse primer 5’-TCGGGAGATAGTAATTAGCATCCGCC) and subjected to sequence analysis. All of these lines were later regenerated to mature maize plants (Supplementary Table 1).

### Library preparation and sequencing

DNA was extracted by the CTAB method ^68^ and libraries were prepared using KAPA Hyper Prep kits and KAPA Single-Indexed Adapter kits for Illumina Platforms (KAPA Biosciences, KK8504 and KK8700). For the lambda experiments, 14 rice and 10 maize lines were skim sequenced at low coverage (~1x) using Illumina NextSeq PE35. Of those, eight rice and four maize lines were chosen for deeper sequencing using Illumina NextSeq PE75, achieving an average coverage of 20X for rice and 15X for maize. For plasmid experiments, six lines each transformed with either pANIC10A-OsFPGS1 or pANIC12A-OsFPGS1 were sequenced with Illumina NextSeq PE75 at ~20X.

### Copy number calculation

The lambda and plasmid sequences were added to the rice ^69^ and maize reference genomes ^70^ as separate chromosomes to construct concatenated genomes, which were then used as references for read alignment by BWA-mem (version 0.7.15) with default parameters ^71^. For skim-sequenced lines, the mean coverage of lambda/plasmid and genome in each event was estimated as the division of the the total number of reads mapped to individual sequences by their respective genome sizes. For lines sequenced at high coverage, the average read depth of lambda/plasmid and genome was calculated as the mean of per-base coverage analyzed by bedtools (version 2.26). The copy number of lambda/plasmid was then derived by multiplying the mean coverage by two, considering that the insertions are heterozygous in diploid genomes.

### SV calling

After adapter trimming by trimgalore (version 0.4.4) and quality checking by fastqc (version 0.11.3) at default settings, reads were aligned to the rice/maize concatenated genomes where lambda and plasmid sequences were added as separate chromosomes, using BWA-MEM (version 0.7.15) with default parameters. PCR duplicates were removed by Picard’s MarkDuplicates (version 2.4.1) and MAPQ filter of 20 was applied. The output BAM files were analyzed for structural variants by SVDetect (version 0.7) ^31^ and Lumpy (version 0.2.13) ^32^ to call inter-chromosomal translocations and intra-chromosomal translocations. For SVDetect, step length and window size were calculated separately for each sample and structural variants supported by fewer than two reads were filtered. For Lumpy, the mean and standard deviation of insert sizes were calculated for each sample, with two reads set as minimum weight for a call and trim threshold set as 0. For intra-chromosomal translocations, the read cutoff for both Lumpy and SVDetect was set at 3 to increase accuracy. Structural variants in each event called (from both Lumpy and SVDetect) were filtered against those of wild type and the other events with an in-house script. Unique breakpoints were manually inspected with IGV (version 2.3.81) and plotted with Circos (version 0.69) ^72^.

### Data simulation

We performed four sets of simulations by embedding shattered and reshuffled fragments from lambda or other chromosomes into the rice reference chromosome 1 sequence at random sites (Supplementary Table 2). Subsequently, a heterozygous diploid genome was constructed by concatenating the modified chromosome with reference chromosome 1. Paired-end Illumina reads were then simulated by ART (version 2.5.8) ^73^ at coverage 10X. For ART, the Illumina sequencing system was set as NextSeq 500 v2 (75 bp), average fragment size and standard deviation were set to 300 and 80 bp respectively. The inter- and intra-chromosomal translocations in each simulated data set were then identified with the SV calling pipeline described above. The output of Lumpy and SVDetect were compared with the simulated data to assess detection performance.

### Junction assembly and validation by PCR

Reads that support lambda-genome junctions identified by both SVDetect and Lumpy were assembled by SPAdes (version 3.10.0) ^74^ with default parameters. The output sequences were aligned against to the reference genome with NCBI blast (version 2.2.26) at default parameters and used as templates for primer design. The blast output of all rice events transformed with lambda were then subjected to analyzing microhomology at junction sites and identifying relative orientations between ligated fragments with an in-house script. Selected products of PCR were sequenced and aligned to the assemblies.

### Copy number variation detection

CNVnator (version 0.3.3) ^33^ was employed to call copy number variation on BAM files where mapping quality was set to be at least 20. The bin sizes for rice and maize genomes were set at 500 and 5,000 bp respectively. The CNVnator output was filtered by removing calls with q0 > 0.5 and eval1 > 0.01 using an in-house script. We declared copy number variation as a deletion if the copy number in specific sample is between 0.5 and 1.5, and at least 0.5 lower than that in wild type and all other samples (unaltered regions are expected to have copy numbers between 1.5 to 2.5). We declared copy number variation as duplication if the copy number in specific sample is between 2.5 and 10, and at least 0.5 higher than in wild type and all other samples. Copy number variations in non-repetitive regions where breakpoints were identified were further inspected using IGV.

### Bionano optical mapping

High molecular weight DNA was prepared from rice λ-4 young leaf tissue using the IrysPrep Plant Tissue DNA Isolation Kit (RE-014-05) and labeled with Nt.BspQI using the IrysPrep NRLS labeling kit (RE-012-10). Data were collected on a single BioNano IrysChip at 80X coverage with an average molecule length of 248 kb. The raw data were assembled with IrysView software set to “optArgument_human”, resulting in 501 BioNano genome maps with an N50 of 1.050 Mbp. The genome maps were then aligned to an silico-digested BspQI cmap of the rice Nipponbare reference genome. Overall alignment was excellent, yielding a “Total Unique Aligned Len / Ref Len” value of 0.946, which exceeds the general recommendation of 0.85 ^34^ Potential SVs were identified using Bionano Solve software (version 3.0.1) and analyzed individually by eye. In addition to the large insertion on chromosome 2, the SV calling software identified several other regions with small (<100 kb), potential insertions in the rice λ-4 sample relative to the Nipponbare reference (Chr3: 31,097,744, Chr12: 20,548,710, Chr1: 2,287,999, Chr3: 13,461,815, Chr3: 16,548,253, Chr6: 3,287,514, Chr7: 22,897,940). As none of these correspond to locations of lambda or plasmid insertions by sequence analysis, they may either represent differences between the Taipei 309 line (used for transformation) and the Nipponbare reference, errors in either assembly, or small insertions caused by biolistic transformation but not involving lambda or the plasmid.

### PacBio sequencing and analysis

High molecular weight DNA was prepared using a modified CTAB method ^75^ from young leaf tissue. The single plant was one of the 23 progeny from the original λ-4 transformant and was homozygous for the large insertion on chromosome 2. The PacBio library was prepared following SMRTbell library guidelines. The library was sequenced with three SMRT cells to generating 10.32 Gb of long reads with N50s ranging from 16-18 kb. Consensus sequences were generated from subread BAM files using Iso-Seq using SMRTLink (version 5.1) with parameters: min_length 50, max_length 30,000, minPredictedAccuracy 0.8, minZScore −3.4, minPasses 0, maxDropFraction 0.34, and polish. The derived consensus reads were then mapped to lambda sequence with BLASR ^76^ at default settings. The BAM output file of mapped reads was converted to fasta format by samtools (version 1.3.1) and aligned against lambda full sequence using NCBI blast (version 2.2.26) at default parameters. The blast output was filtered by removing reads with E-values higher than 0.1 and lengths shorter than 30 bp, and by retaining the longest consecutive matches for each read using an in-house script.

## ACKNOWLEDGEMENTS

We would like to thank Peter Lafayette, Melissa McGranahan, and Gary Orr for help with rice biolistic transformation. We appreciate the help and guidance of Jonathan Gent and Magdy Alabady in bioinformatic analyses. We would like to thank Jean-Michel Michno and Jonathan Gent for critically reading the manuscript.

## SUPPLEMENTARY FIGURE LEGENDS

**Supplementary Fig. 1 | Circos plots of additional rice lines transformed with λ and plasmid pPvUbi2H**. For rice event λ-6, the coverage of λ is divided by 8.

**Supplementary Fig. 2 | Three major intra-chromosomal SV types and the strand orientations of paired-end reads**. **a**, Deletion-type intra-chromosomal rearrangement. b, Duplication-type intra-chromosomal rearrangement. **c**, Two types of intra-chromosomal translocation-type rearrangements.

**Supplementary Fig. 3 | Additional data from maize lines transformed with λ and plasmid pBAR184**. For maize event λ-2, the coverage of λ and plasmid are divided by 10 and 8 respectively. **b**, Swarm and violin-plots showing deletions, duplications and triplications in all maize events transformed with λ. Each dot in the swarm plots represent a different SV. Violin plots represent the statistical distribution, where the width shows the probability of given SV lengths.

**Supplementary Fig. 4 | Linkage analyses of fragments from the 1.6 Mb array of rice λ-4 in self pollinated progeny. a**, Schematic diagram of three genomic fragments flanked by lambda pieces embedded within the 1.6 Mb array on chromosome 2. Two of the fragments are from chromosome 9 (colored black), of size 102 bp and 464 bp respectively, while the other is a 108 bp segment from chromosome 12 (colored red). The arrows indicate the 3’ ends of λ and genomic fragments. The positions of primers P1 to P10 for five amplicons are indicated. **b**, Three fragments from chromosome 9 and 12 are genetically linked with the insertion borders on chromosome 2. Among the 23 T1 plants, 12 are heterozygous for the insertion on chromosome 2 (blue), 5 are wild type (grey), and 6 are homozygous for the insertion (orange).

**Supplementary Fig. 5 | Distributions of microhomology at junction sites and relative orientations of rejoined fragments. a**, Sequence of the 1.1 kb rearranged fragment, where the color code matches that in Figure 2b. Regions of microhomology at the breakpoint junctions are highlighted in cyan, and novel insertions are highlighted in magenta. **b**, Distribution of blunt-ends, insertion of novel sequence, and microhomology of 1-4 bp, 5-25 bp and >25 bp at breakpoint junctions. **c**, Distribution of four relative orientations of rejoined fragments, HH (head-head), HT (head-tail), TT (tail-tail), TH (tail-head).

**Supplementary Fig. 6 | Circos plots of additional rice lines transformed with plasmid pANIC10A-OsFPGS1**. The coverage of plasmid 10A is divided by 2 in 10A-1 and 10A-4, and by 5 in 10A-2.

**Supplementary Fig. 7 | Circos plots of additional rice lines transformed with plasmid pANIC12A-OsFPGS1. b**, Copy number pattern of the highlighted region on chromosome 8 in 12A-3 is exhibited in lower panel. **c**, Swarm and violin-plots showing the distribution of the size and number of deletions, duplications and triplications in all events transformed with single plasmids 10A and 12A. Each dot in the swarm plots represent a different SV. Violin plots represent the statistical distribution, where the width shows the probability of given SV lengths.

